# Reconciling Flexibility and Efficiency: Medial Entorhinal Cortex Represents a Compositional Cognitive Map

**DOI:** 10.1101/2024.05.16.594459

**Authors:** Payam Piray, Nathaniel D Daw

## Abstract

The influential concept of a cognitive map envisions that the brain builds mental representations of objects, barriers, and goals. This idea has been formalized in a range of computational models that show how such representations can be useful for guiding goal-directed behavior, for instance by planning novel routes to maximize long-run rewards. One key feature of flexible cognitive representations generally is that they exploit compositionality – the ability to build complex structures by recombining simpler parts. However, how this principle plays out in neural representations of cognitive maps and map-based planning remains largely unexplored. Indeed, as we show here, compositionality can be difficult to reconcile with efficient planning: because reuse tends to oppose flexibility, it is challenging to construct a compositional representation of the environment which is also organized in a way that enables generalizability and efficient planning. Here, we propose a novel model for efficiently creating and planning with compositional predictive maps, and further show that it successfully simulates various aspects of response fields in the medial entorhinal cortex, particularly object vector cells and grid cells. The model treats each object as an alteration to a baseline map linked to open space, creating complete predictive maps by combining object-related representations compositionally. Overall, this work provides a comprehensive and realistic model for efficient model learning and model-based planning in animals, and offers insights into the brain processes supporting efficient, flexible planning using compositional predictive maps.

## Introduction

The principle of compositionality – the ability to reuse representations constructively to build up novel, complex ideas from simpler parts – is widely viewed as fundamental to intelligence. In cognitive science and neuroscience, it is thought that the brain’s impressive capacities for perception, language, and reasoning arise, at least partly, from its compositional nature, assembling thoughts and concepts from simpler components (Bienenstock et al., 1996; Chang et al., 2017; Frankland & Greene, 2020; Lake et al., 2017; Spelke, 2022). Similarly, in artificial intelligence, compositionality is seen as an important capability for developing human-level artificial general intelligence. While many modern artificial intelligence systems do not yet exhibit human-like compositional abilities (Fodor & Pylyshyn, 1988; Gershman et al., 2015; Goyal & Bengio, 2022; Hill et al., 2017; Lake et al., 2017), creating models that demonstrate compositional behavior has been a central goal driving advances in the field (Greff et al., 2020; Lake, 2019; Lake & Baroni, 2023; Liu et al., 2023; Meng et al., 2022; Nie et al., 2021).

One case in which structural learning has been studied in detail is that of trial-and-error action selection, particularly reward maximization in sequential tasks such as spatial mazes. It has long been argued that animals accomplish this, at least in part, by learning a “cognitive map” or internal representation of the environment and task (Behrens et al., 2018; McNaughton et al., 2006; O’Keefe & Nadel, 1978; Schacter et al., 2007; Tolman, 1948). This is thought to enable them, for instance, to plan new routes to novel goals and find detours and shortcuts. In turn, these functions have been widely formalized computationally using a family of algorithms from reinforcement learning (RL), known as model-based learning because they rely on a learned internal model of the environment’s contingencies.

Interestingly, however, although such map or model learning also might seem to involve the composition of meaningful subparts (e.g., a building interior is made up of hallways, rooms, and doors) and this has occasionally been suggested (Ho et al., 2022; Hunt et al., 2021; Tsividis et al., 2021), this aspect has not been widely studied or theoretically developed. Instead, map learning in the RL setting has most often been formally characterized as akin to individually discovering and memorizing each of the edges in a graph (Daw et al., 2005; Glascher et al., 2010). Here, we suggest that this gap reflects a key challenge in constructing and using such maps: creating representations that balance competing demands of flexibility vs. efficiency. In general, using a (“one-step”) map or model to flexibly plan new paths to goals requires significant computation to search iteratively over candidate paths. Meanwhile, optimizations that address this by reusing or “caching” some of this work (for instance, chunking useful extended action sequences) generally simplify computation at the expense of reducing generalizability to new situations (e.g., where different action sequences would be better). Exploiting compositionality in this setting thus remains a challenge, because representations that cache demanding computations for efficient planning resist flexible, compositional construction. Accordingly, to the extent a few previous theories of cognitive maps have captured some compositional-like relational structure in the task graph (Behrens et al., 2018; George et al., 2021; Whittington et al., 2020), these models have not generally addressed using these maps to plan or seek reward and addressing this would require exhaustive and arguably implausible computation at decision time.

Here we introduce compositionality into on an alternative class of RL models (known as successor representation models) which have recently become popular for enabling efficient planning, by representing the cognitive map in terms of long-range predictive dependencies across the task space rather than by local adjacency (de Cothi & Barry, 2020; Piray & Daw, 2021; Rueckemann et al., 2021; Stachenfeld et al., 2017). In particular, we build on a recent development of this class (the “default representation”; (Piray & Daw, 2021)) that, by introducing a particular nonlinearity and some linear algebraic tricks, largely addresses the issues of generalizability and flexibility. Here we extend this model to address two types of open issues in this domain. First, computationally, we introduce compositionality so as to address how such predictive maps can be built quickly but also in a generalizable way by recombining precomputed, reusable representations. Second, neurally, we argue that a number of aspects about how this model decomposes maps into stable and reusable pieces shed new light on observed neuronal firing patterns in the medial entorhinal cortex (MEC), which previous modelers have suggested may subserve a neural representation of maps of this sort (Stachenfeld et al., 2017). Specifically, we argue previous models fail to account for important aspects of cellular responses in the MEC showing composition-like representations, such as invariant and modular coding response in MEC object vector cells, and how these coexist with fast and stable coding in MEC grid cells.

More specifically, we propose a novel model for constructing compositional predictive maps that can use the resulting maps to perform complex planning tasks efficiently and flexibly. The presented model is based on the idea that each object can be seen as an alteration to a baseline predictive map of an open-field space. For each object, we derive a representation of its contribution to the predictive map that does not depend on the specific position or rotation of the object in the task space. The complete predictive map of a state space consisting of multiple objects is then built by putting together the representations related to each object. The resulting map can then be used directly and efficiently for planning, without laborious search.

With regard to the function of the MEC, the model suggests that different spatially tuned cells within the structure can be understood as encoding components of the compositional predictive map. In particular, the theory accounts for a class of cells discovered recently in the MEC, object vector cells, which fire exclusively when animals are near objects (Andersson et al., 2021; Høydal et al., 2019; Kinkhabwala et al., 2020). These cells are the most ubiquitous class of cells in the MEC, accounting for roughly 30% of all spatially tuned cells in this region (Høydal et al., 2019). According to our model, these cells encode the building blocks of compositionality, a planning-ready computational representation of objects, borders and other components of the map specialized in a planning-ready way that enables the system to flexibly build a map and easily use it for planning. In turn, our theory interprets grid cells as providing an efficient code that provides a baseline map over which these components can be incorporated. We show that this successfully models various empirical findings concerning the firing patterns of grid cells following the introduction of barriers and goals.

## Results

### Default Representation (DR)

We build on our recent computational model of map learning and planning in the brain, linear RL (Piray & Daw, 2021), which exploits long-range predictive maps for planning and shows animal-like behavior across various replanning tasks, such as reward revaluation (e.g., Tolman’s latent learning task) or transition revaluation (e.g., Tolman’s detour task).

To briefly summarize (see Methods and (Piray & Daw, 2021)), as with other RL theories, linear RL formalizes planning as computing the value function, or the expected sum of future rewards for different possible courses of action. More specifically, the value function *v*^*^(*s*) captures the sum of rewards (minus costs) expected by starting at state (e.g., location) *s* and following an optimal trajectory thereafter. This function contains sufficient information to choose optimally (e.g., by visiting the highest-valued neighboring location at each step). Classic RL algorithms for model-based planning start by learning the one-step adjacency map of the environment, **T** (a matrix of dimension *N* states by *N* states representing transition probability between states under a random walk), together with a vector of rewards or costs in goal states, **r**, and using these to iteratively sum rewards over candidate trajectories.

The key idea of linear RL, like the successor representation (SR) is to simplify this laborious iterative procedure by computing from the one-step adjacency map **T** a long-range predictive map **D** = (diag(exp(**c**)) − **T**)^−1^, called the DR (see more detailed definitions in Methods). Here diag(exp(**c**)) is a diagonal matrix whose diagonal values depends on per-state costs vector **c**. The DR in effect captures, for each state, which other states will be visited in the long run following it, and what cumulative per-step costs **c** will be expended reaching them. Given **D**, it is possible to compute the value function *v*^*^ without extensive iteration, but instead by a single matrix-vector multiplication, which is easily implemented in a single layer neural network. (Here, the vector depends on goal rewards **r**, which explains how the system can easily adapt to changes in goals.) A main intuition is that the matrix inversion in the definition of **D** arises algebraically from and is equivalent to iteratively summing costs over many transition steps (assuming that the transition is done under the default policy); once this computation is done, planning is easy. Thus, in the present work we solely focus on the computation of the DR map **D**, and in particular on exploiting compositionality to flexibly compute the matrix inversion by reusing precomputed components.

### Building the DR compositionally

We present a computational model for making predictive maps that builds the DR by putting together representations corresponding to different objects in the environment. Formally, these objects correspond to stereotyped configurations of barriers (e.g., rooms, shapes, or walls; Fig. 1) that interrupt travel in the open field. In this case, each object can be seen as a perturbation of the open field. Assuming we start by knowing as baseline the DR for the open field, **D**_*os*_, then matrix algebra (specifically, the Woodbury matrix inversion lemma) implies that the DR for an environment containing a single object can be written by adding a correction term to **D**_*os*_:

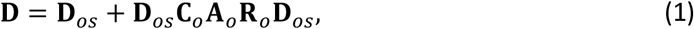

**Figure 1.**
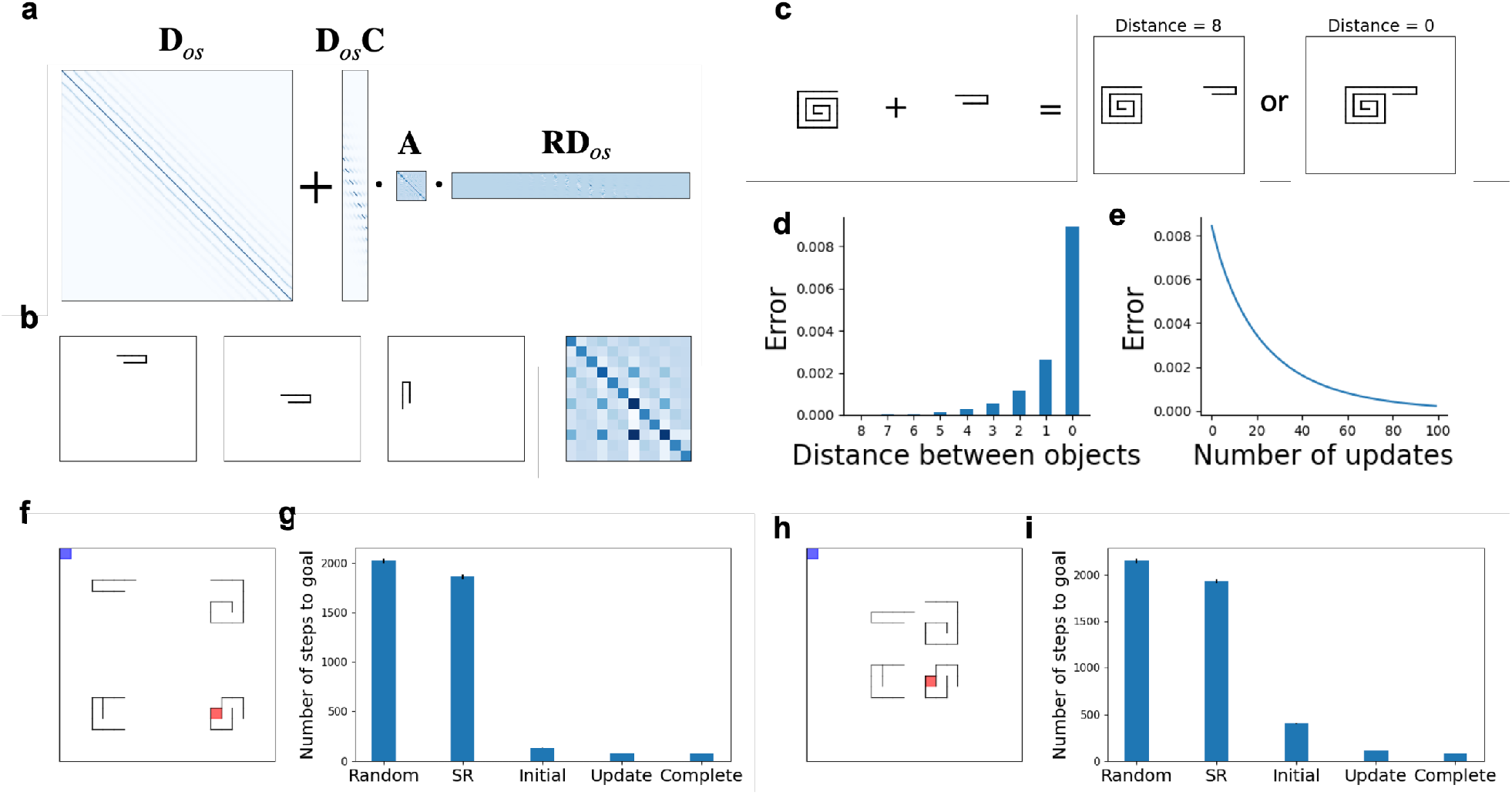
The compositional predictive map enables efficient planning. **a**. The model represents any object as a perturbation of the open field. For computing the predictive map when a single object is introduced, the only nontrivial computation is related to computing the predictive object representation matrix, *A*, which its size depends on size of the object rather than the entire state space. **b**. The predictive object representation matrix is invariant to translation and rotation. Here, matrix **A** is displayed for an object, its translation and rotation. **c, d**. Two objects are added to create a new environment as a function of distance between the two objects. Their corresponding predictive object representation matrix provides a first order approximation for the new environment. The error between this approximation and the true predictive object representation matrix is plotted as a function of distance between objects. The error increases as the objects get closer together. **e**. The error function is plotted for the learning algorithm that learns the true predictive object representation matrix for the environment with the two objects plotted in **c** (the one with Distance=0). With each update, the error decreases. The starting point of the algorithm is given by putting together the corresponding matrix **A** of the two objects. **f**. An example planning environment, in which blue and red squares indicate start and terminal point, respectively. **g**. The predictive map for a new environment was constructed using our compositional approach and a number of alternatives. This involved either adding the PORs for the four objects (referred to as the ‘Initial’ model), or utilizing an ‘Update’ model that also updates the initial map once per every step using the algorithm introduced above. Additionally, a ‘Complete’ model, which builds the predictive map entirely, was considered. This model is an extension of the update model, in which the update process is continued until convergence. Furthermore, we simulated the successor representation (referred to as ‘SR’) to create predictive maps, which updates the maps using a temporal difference algorithm. Finally, we considered a random walk algorithm (referred to as the ‘Random’). Overall, both the Initial and Update models greatly outperformed the random walk and the SR, and it was on par with the performance of the Complete model. **h, i**. A more challenging environment in which objects are closer to each other. As expected, the Initial model was performed worse, although its performance was still comparable to the Complete model. Simulations in f-i were repeated 5000 times and errorbars (might not be visible) represent standard error of the mean.

While this expression may appear notationally complicated, it is computationally quite simple, and much easier and more plausible than re-inverting the whole matrix from scratch (Fig. 1a). First, it expresses a compositional construction, in which the map is expressed as the sum of the baseline map plus the object-dependent perturbation, reflecting the ways in which that object affects predicted long-run trajectories between locations in the environment. Although each of the summed terms is a matrix of size *N* × *N* (where *N* is the number of locations in the environment), the perturbation matrix is simple in a formal sense, i.e., low rank.

Specifically, the only nontrivial computational term in this equation is the matrix **A**_*o*_. We call this matrix the predictive object representation (POR) for the corresponding object. Importantly, the dimensionality of the POR depends on the size of the object, not the whole state space (Fig. 1a). This is because introducing an object to an open space changes one-step transitions locally (i.e., in the vicinity of the object) and its consequences for other locations across the long-range predictive map, **D**, can then be computed efficiently using matrix algebra (see Methods). This is accomplished by the other matrices which index these effects: **C**_*o*_ is a binary column matrix whose columns encode the location of states that are immediately next to the object, and **R**_*o*_ is a row matrix that is nonzero only in states that are immediately next to the object. In other words, matrices **C**_*o*_ and **R**_*o*_ serve as simple lookup tables encoding the one-step relations of the object with other states. Their product with the baseline map **D**_*os*_ then gives a linear combination of either columns or rows of **D**_*os*_ that are immediately related to the object, a computation that can be readily implemented in a simple neural network with fixed, pre-initialized weights.

An important insight is that the POR is translation- and rotation-invariant, i.e., it does not change if one moves or rotates the object in the state space (Fig. 1b): the effect of any object on trajectories is clearly invariant up to the frame of reference, and the **C**_*o*_ and **R**_*o*_ matrices align the object in the broader map.

It is also important to stress what this procedure accomplishes. In classic model-based RL, it would of course be straightforward to derive the one-step transition map **T** compositionally by directly adding a set of barriers to an open grid. However, following any such update, it would be necessary to recompute the optimal values by a full matrix inversion (to produce **D**), or alternatively, summing over a large number of trajectories. Here, instead, we compositionally build the long-run predictive map **D** directly, reusing most of the computation underlying the matrix inversion via **D**_*os*_ and **A**_*o*_ (the learning of which we address below). Updated optimal action values (and the optimal policy) can then immediately be produced with a matrix-vector multiplication.

To construct the DR for environments with multiple objects, we can, to a first approximation, simply sum PORs from multiple objects. This is because if two or more objects are far from one another, their combined POR is approximately equal to the block-concatenation of their corresponding PORs. Thus, formally, if **A**_1_ and **A**_2_ are PORs of such two objects, the predictive DR map is approximately given by:

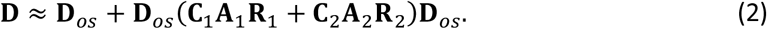

In this way, a set of PORs can be viewed as a set of basis functions for predictive maps reflecting common object motifs in a set of environments. This in turn could straightforwardly provide the foundation for a simple learning procedure (not elaborated here) for exploring a new environment and quickly estimating its corresponding map **D**, while also efficiently planning at each step of learning.

Note that the convenient compositional expression from Eq. 2 is simplified with respect to the exact DR. (The latter would be more laboriously obtained by the iterative application of Eq. 1, such that the perturbed **D** replaces **D**_*os*_ when adding **A**_2_ and so on, and thus the object effects do not decompose into a simple sum.) The intuition for the difference (Fig. 1c) is that when two objects are close (and especially when they touch) their effects on possible trajectories through the environment can be different from those implied by either object alone. Nevertheless, even when objects are close, this equation provides a first-order approximation of the POR of both objects (Fig. 1de).

Moreover, we have developed an efficient and biologically plausible algorithm (see Methods) to iteratively update the combined POR for two or more objects and learn it gradually (either by experience or mental replay or both), correcting the initial approximation. The same algorithm is applicable to the case of learning a single POR like **A**_1_ from more elemental parts (corresponding ultimately to individual barriers). This algorithm is computationally efficient because it works in the space of objects (i.e., the part of the space containing the subcomponents: the size of the combined POR), rather than the whole state space (the size of **D**). This is crucial because previous algorithms for learning predictive maps by experience, notably the SR (Dayan, 1993; Gershman, 2018; Momennejad et al., 2017; Russek et al., 2017), were not scalable, because they had to update the entire predictive map matrix in each iteration. In practice, the proposed algorithm converges quickly, usually within few trials, which is mainly because Eq. 2 already provides a good approximation of the combined POR (Fig. 1f).

In all, this learning procedure describes how a set of PORs can be built up for stereotyped objects like rooms and, in turn, how the combined PORs from multiple objects close to one another in a new environment can, with learning, be adjusted to a single corrected POR for their combination. This could, in turn, provide the basis for a metalearning algorithm (which we do not pursue here) that learns an appropriate basis set of PORs matched to a set of object motifs reoccurring across a family of environments.

### Planning with a compositional DR

This approach enables efficient planning. To demonstrate this, we conducted a simulation analysis in which different models were exposed to four different environments, each with a distinct object. This allowed the model to learn four PORs. Subsequently, models were simulated in a new environment containing all four objects. For comparison, we considered the random walk as well as the SR, an alternative way to construct predictive maps experientially (non-compositionally), based on a temporal-difference learning rule (Dayan, 1993; Russek et al., 2017; Stachenfeld et al., 2017). We built the predictive map in two different ways and compared their planning capabilities with these baseline models, as well as a complete predictive map.

The first approach involved a simple first-order approximation model, as described above, where PORs for all objects were simply added as in Eq. 2. We used this map with the linear RL algorithm, enabling us to plan a route from a starting point to a terminal point. The second approach additionally updated the combined POR for the four objects using the learning algorithm mentioned above. We assumed that the time required for an update step was equal to the time needed for taking a real navigational step in the world, although it’s worth noting that the updates can often be much faster.

As depicted in Fig. 1f-g, even the first model, which was simply built by a linear composition of individual PORs, performed relatively well. This is important because the computational load of this algorithm is nearly as low as that of a random walk. However, unlike the random walk, which rarely reaches the target in a reasonable time, this algorithm navigates to the target effectively (with an average path length compared to the ground-truth planner of 1.76). Moreover, even with a slow update rate (i.e., one update per step), this algorithm achieved accuracy close to the ground-truth planner (with a relative average path length of 1.09). Importantly, as described above, the update is in the space of objects, not the whole state-space, which makes this algorithm much more efficient than previous algorithms, such as the SR, for creating a predictive map. Additionally, it is worth noting that the ground-truth model can be seen as an asymptotic case of the update model, after updates have been performed until convergence.

We repeated this analysis under more challenging conditions, where the objects were placed closer to each other. As expected, the performance of both algorithms, especially the Initial model, declined, but remained relatively comparable to the complete planner and substantially better than the random walk and the SR (Figure 1h-i).

### Neural representations of map components in the MEC: Object vector cells

A number of previous modeling studies have suggested that spatially tuned neurons in the MEC may in some way collectively represent a cognitive map. For instance, Stachenfeld et al. suggested that grid cells of MEC collectively resemble a factorization of the environment’s SR, resembling its eigenvectors (Stachenfeld et al., 2017). A second recent proposal, known as the Tolman-Eichenbaum Machine (Whittington et al., 2020) develops a related proposal more implicitly by training a network, via backpropagation, to learn motifs of graph connectivity across many environments (similar, in our terms, to meta-learning the one-step map **T**). Units inside the trained network have responses resembling those of several types of spatially tuned MEC neurons. The present model elaborates these accounts and, through more explicit analysis of the mapping and planning problem, clarifies the computational mechanisms that, we suggest, underly these observed correspondences.

Recent research in the MEC reports a novel family of neurons known as object vector cells (Høydal et al., 2019). These neurons activate when an animal is at a certain distance and direction from an object. It has been observed that 20-30% of the neurons in the MEC are object vector cells, which makes them the most common cell type in the MEC, well above grid cells, head direction cells, and border cells.

We suggest these cells can be understood as capturing the perturbation of the map due to the inclusion of the object, i.e. the POR term from Eq. 1. Perhaps the most significant characteristic of object vector cells is their translation invariance. In a study by Høydal et al. (2019), MEC cells’ response fields were examined in two consecutive trials. An object was initially placed on the floor and then moved to a new location. The researchers observed that object vector cells that fired initially exhibited a similar response even after the object’s translation. Importantly, the displacement from the object was preserved. Similarly, the model, i.e. the projection of the POR into the state space, demonstrated a consistent response across both trials due to its translation invariance (Fig. 2).

**Figure 2.**
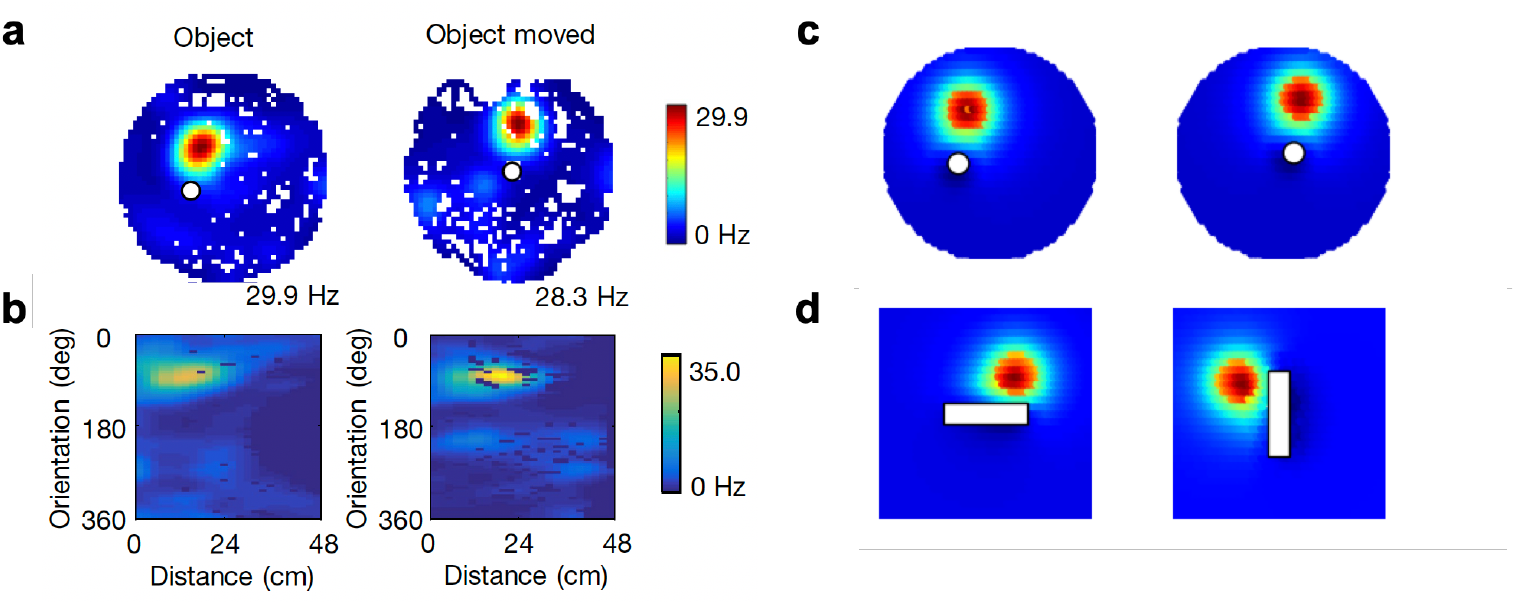
Object representations are invariant to distance and orientation relative to the object. a) Firing rate maps of an example object vector cell reported by Hoydal et al. When the object was moved, the firing field was moved accordingly. b) The vector map for the cell shown in a, as a function of allocentric orientation and distance from the center of the object. c-d) model simulations show similar properties when the object was moved (c) or rotated (d).

Furthermore, Høydal et al. (2019) observed that MEC object vector cells exhibit a consistent response to an object when the animal is at a similar distance and orientation relative to the object. The model displays a similar behavior, as the PORs are rotationally invariant (Fig. 2).

Note that we model object vector cells as a row vector of the POR corresponding to the specific state associated with the particular distance and direction from the object to which the cell is tuned. Generally, this formulation provides a way to account for the activity of “vector” cells in the MEC (Bicanski & Burgess, 2020), such as object vector cells or boundary vector cells, within the RL framework that assigns a unique state to each spatial location. Vector cells in the MEC exhibit selective firing patterns exclusively when the animal is positioned at a particular distance and direction from the cell’s focal point, which could be an object or a boundary in the environment. Within the RL framework, this firing behavior is effectively equivalent to the activity pattern corresponding to the state that shares the same directional and distance relationship with the object or boundary as the animal’s current location. Object vector cells serve as an intermediary step in the process of constructing the final representation of space in the form of grid map (which we cover later), and their object-relative code facilitates and reflects the compositional construction of the final grid map.

Another property of object vector cells is their ability to generalize across different objects and environments. Unlike landmark vector cells in hippocampal area (Deshmukh & Knierim, 2013), object vector cells do not differentiate between different objects as long as they introduce similar disruption to the state space. The model also shows the same behavior. Since PORs capture perturbation of trajectories through the state space by the object, not its specific properties, they remain the same as long as the way that the object impedes passage remains the same. Furthermore, empirical data showed that object vector cells, unlike landmark vector cells, do not depend on experience and emerge almost from the first trial, even in a novel environment (Fig. 3). The model shows the same behavior because PORs are not learning or experience dependent (unlike predictions of the model for grid cells as shown in the next section.) Furthermore, when multiple objects when introduced in the field, object vector cells fire in response to each with small differences in distance and orientation (Fig. 3). The model accounts for this because PORs do not encode interactions between objects, and therefore we expect them to generalize to multi-objects environments.

**Figure 3.**
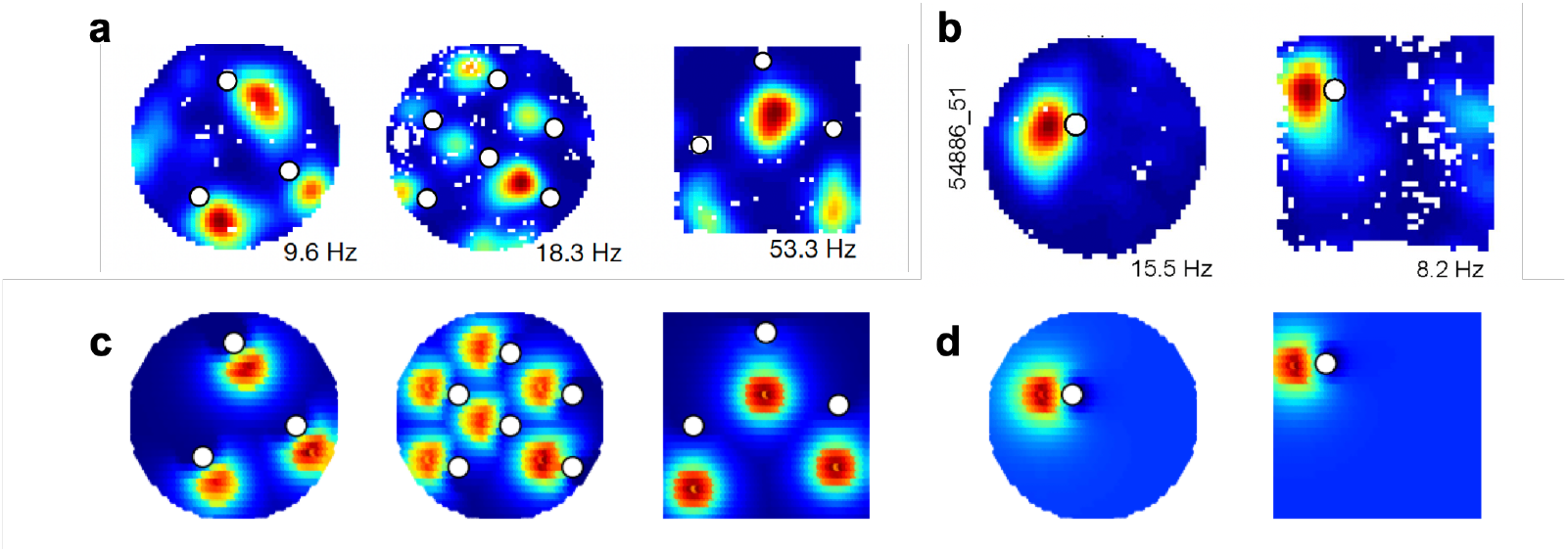
Object representations generalize to multi-object environments, and to novel objects and novel environments. **a**. The firing field for three object vector cells in a multi-object environment in which a few new or old objects (with similar sizes) that have been placed in the space. Regardless of familiarity, the cells exhibit similar firing fields in the vicinity of each object. **b**. The firing field for an object vector cell in two different environments, which shows that the cell generalizes between environments. **c, d**. Simulation of the model in similar setups. The model shows the same behavior because PORs are only a function of the object size, not its shape or familiarity, and they are not learning or experience dependent. Data in a-b were reported by Hoydal et al.

In their study, Høydal and colleagues found that object length has a significant effect on the firing field of object vector cells, although some variability was observed (Fig. 5). Our model exhibits similar behavior, because the length of an object affects the way it alters state transitions (Fig. 5). However, the model predicts that increasing the height of an object should not affect the state-space since the height does not impact how the object affects trajectories (apart from climbing or jumping). This is also in line with the empirical results reported by Høydal and colleagues.

**Figure 4.**
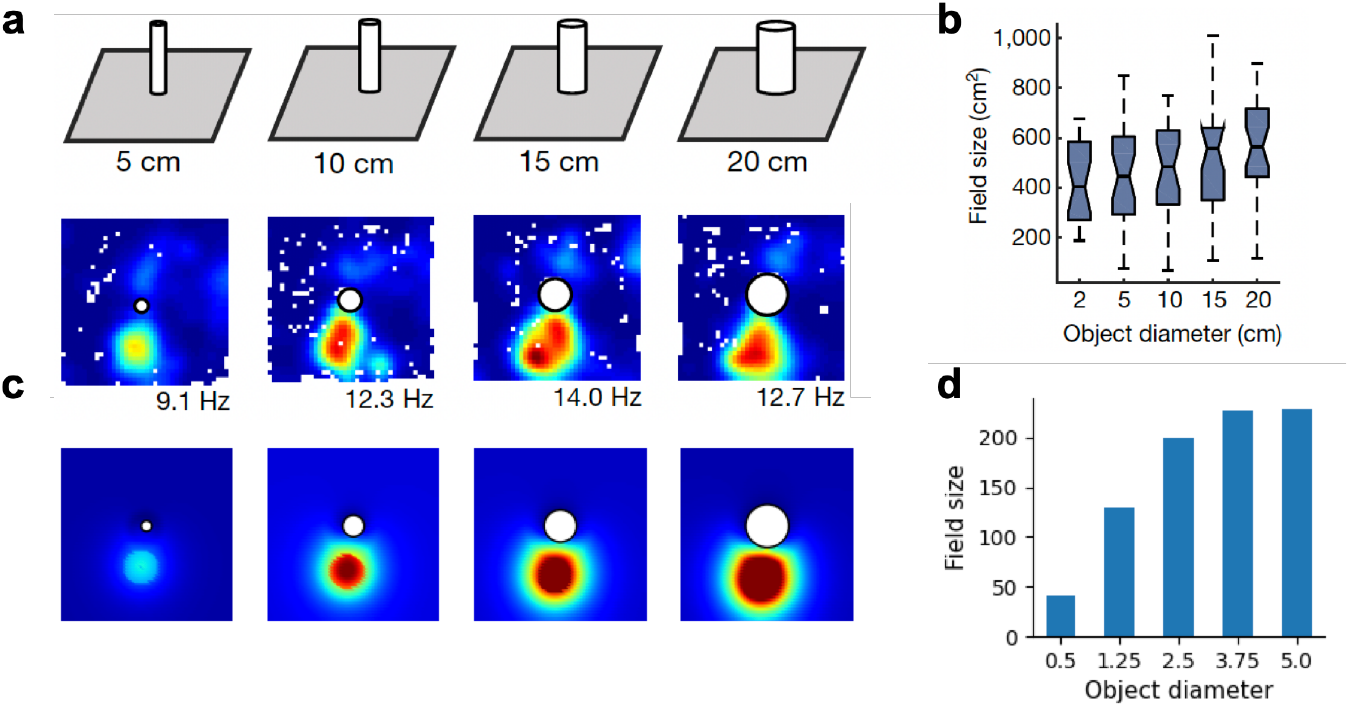
Object representations depend on the object size. **a**. The firing rate for an object vector cell illustrates dependency to the object diameter. **b**. Distribution of field sizes for all object vector cells shows significant dependency to the object diameter. **c, d**. Simulation of experiment results in a-b. Data in a-b is reported by Hoydal et al.

**Figure 5.**
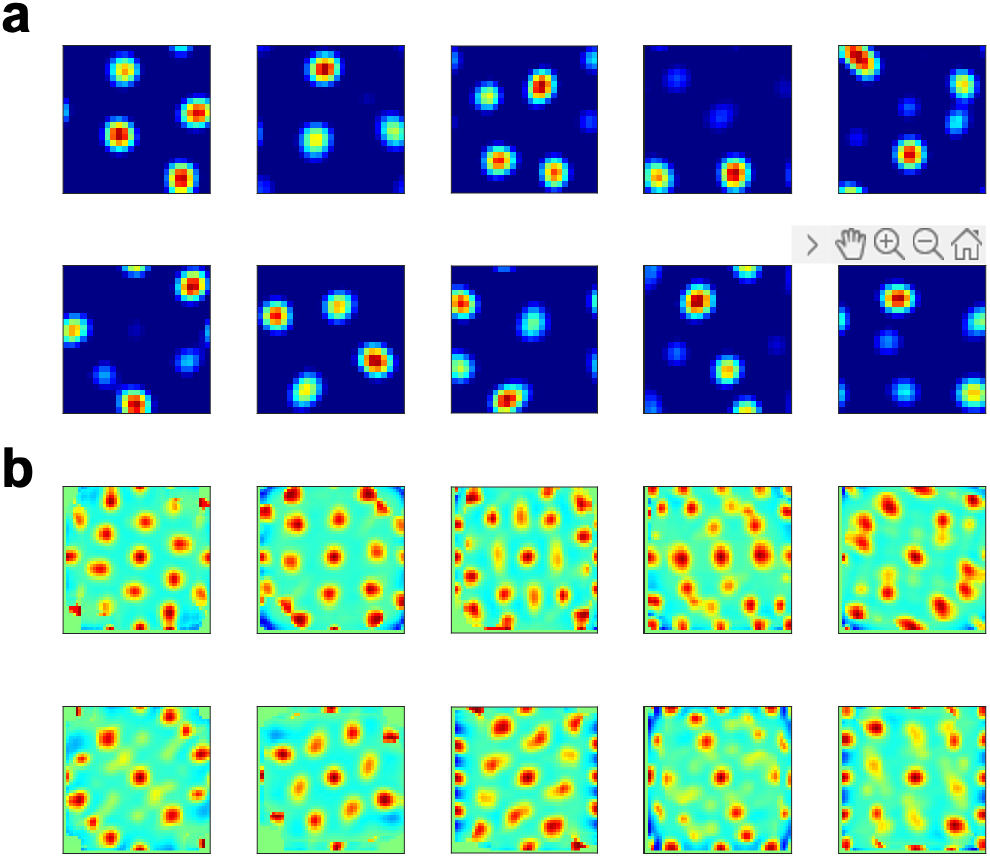
Open space grid code in the model based on Dordek and colleagues. **a**. Example column vectors of **U**_*os*_ unwrapped into the corresponding coordinates of actual space are plotted. The vectors represent a low-dimensional reduction of the open space predictive map. **b**. Spatial autocorrelation maps corresponding to columns are plotted. A substantial number of vector columns of **U**_*os*_ show a grid-like pattern.

A final observation, perhaps less obvious under the current model, is that object vector cells fire even when the object is suspended allowing passage below (Høydal et al., 2019). In the model, this may be because PORs in our model encode computational representations that are precomputed and could be added to the predictive map if needed. These parallel precomputations could allow the system to be flexible and quickly responsive to new situations that might happen, e.g., a suspended object might fall any second.

### Neural representation of the compositionally constructed map: Grid cells

It has been proposed that grid cells encode a low-dimensional decomposition of the predictive map (Stachenfeld et al., 2017). Specifically, in an influential study, Stachenfeld and coauthors proposed that grid cells encode eigenvectors of a different variant of a predictive map, the SR: collectively, these neurons response fields capture the map, in effect as a set of basis functions for it. While this proposal provides a promising characterization of grid cell responses in the open field – particularly when the eigendecomposition is subject to a biologically motivated non-negativity constraint, which improves the match to neural responses (Dordek et al., 2016) – a number of puzzles remain, particularly concerning how the grid code behaves in the presence of barriers. In general, the addition of barriers to the map requires laboriously re-computing the SR (this is the problem addressed by the current model) and its eigendecomposition, and this in turn has substantial effects on all the eigenvectors (i.e., each cell’s simulated tuning). Indeed, even allowing for such recomputation, this model fails to capture the relatively stable code of grid cells when objects and goals are introduced, as eigenvectors tend to change dramatically when the map changes even slightly.

Here, we leverage the current model to propose that grid cells provide a basis space for representation of the DR, which is expanded compositionally to include perturbations due to the PORs as described above. Importantly, the resulting grid code remains globally stable because these perturbations are represented in a space given by the eigenvectors of the open field.

In particular, we consider a baseline model given by the eigenvectors of the DR from the open space, given by the columns of a matrix **U**_*os*_. These are computed by non-negative principal component analysis of **D**_*os*_, using the model of (Dordek et al., 2016), who showed such components can be learned through a single-layer neural network, with Hebbian-like learning rule (Oja, 1982). This procedure produces hexagonal grid patterns (Figure 5).

We then introduce perturbations from PORs as before (presumably arising from object vector cells), projected onto the basis **U**_*os*_. Formally, we propose that grid fields encode the following map:

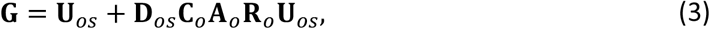

where **G** is the grid code with column as grid cells. Importantly, **G** essentially provides a representation of the DR **D**, as the full predictive map can be easily assembled via the product of **G** with the transpose of **U**_*os*_. Moreover, **G** can be compositionally expanded to incorporate additional objects, analogous to Equation 2. Furthermore, the grid code **G** equals the baseline grid code **U**_*os*_ (and its outer product with itself represents the baseline, open field map **D**_*os*_) when no objects are present in the environment.

Here, we present simulation results examining how different types of barriers and goals in the open space impact the baseline grid code. Thus, we examine the effects on **G** when task-relevant objects are introduced.

A key prediction by our model is that the introduction of barriers and objects should not result in a global remap of grid cells. Instead, the model predicts that such changes in the environment result in local remapping of grid cells. This is because such barriers and objects mainly influence the grid codes near the area of change, through the object-specific lookup table matrices, **C**_*o*_ and **R**_*o*_. This contrasts with the model by Stachenfeld et al., which predicts substantial global grid remapping since barriers alter all eigenvectors (Stachenfeld et al., 2017).

To showcase this, we first consider an experiment by Wernle et al., which examined grid cell changes when environments merge (Wernle et al., 2018). They recorded grid cells in rats exploring two adjacent rectangular compartments, A and B, separated by a dividing wall. When the partition was removed, allowing exploration of the merged compartments, the researchers found grid patterns were largely preserved near the original distal walls (Figure 6). However, approaching the former partition line, individual fields “almost immediately” reorganized to establish spatial continuity between the two maps (Wernle et al., 2018). Specifically, grid periodicity emerged locally around the area where the compartments had met. Thus, their results demonstrate that when merging occurs, grid cells quickly reconfigure to reconcile the separate maps, enabling coherent navigation in the new unified space. The model predicts a similar pattern, because the partition wall primarily influences the area near the partition, due to the relative selectivity coded by object-specific lookup table matrices, **C**_*o*_ and **R**_*o*_.

**Figure 6.**
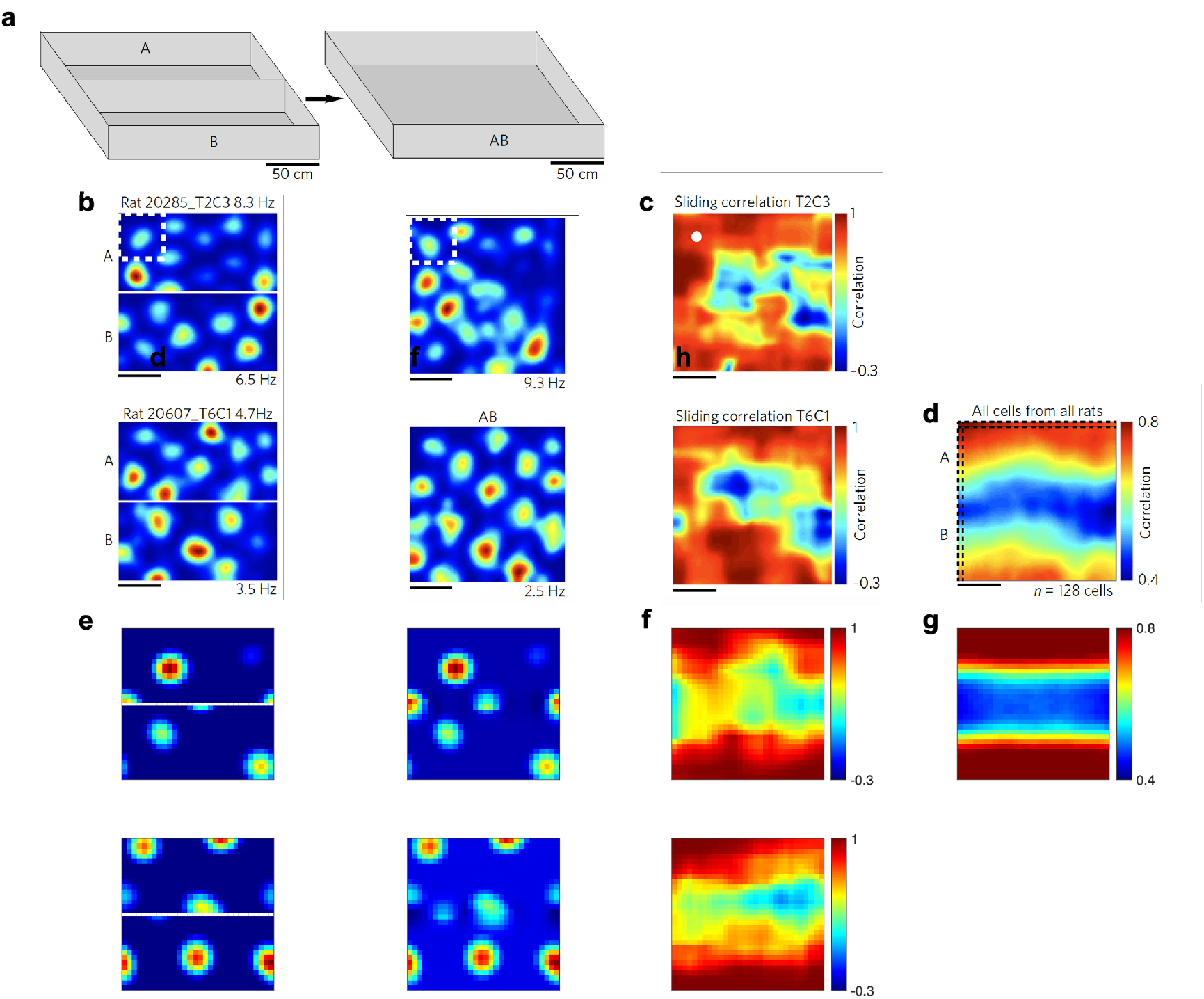
Grid maps change locally after removing a wall. **a**. The experimental procedure of Wernle et al. is illustrated. The rat underwent training in two rectangular compartments, labeled A and B. These compartments were initially separated by a central partition. During the test phase, the partition was removed, allowing the rats to explore the combined open square environment, denoted as AB. **b**. Example firing rate maps for two grid cells from different rats are shown in both the divided A|B and merged AB environments. **c**. The sliding spatial correlation heatmap is shown for the two example cells plotted in b. Spatial correlation is calculated in moving boxes with the size of the white stippled box shown in b. **d**. The average spatial correlation heatmap across all recorded cells is shown. The impact of removing the partition is mostly limited to locations near the partition. **e-g**. Simulation of experimental results in **b-d** by the model. The simulation results in **g** were repeated 50 times with different randomization seeds and the average has been plotted. Data in **b-d** is published by Wernle and colleagues (Wernle et al., 2018).

This property of grid fields was also recently highlighted in another study examining the effects on grid fields of introducing the rat’s home cage into an environment (Sanguinetti-Scheck & Brecht, 2020). In this study, the authors found that grid cells did not globally remap when the home cage was placed in the arena. Instead, they showed that the introduction of the home cage in the center of the arena retained the global representation but resulted in a local shift of grid fields near the embedded home (Figure 7). Thus, the mean normalized rate was selectively increased near the home cage, which is also visible in spatially averaging peak normalized rate maps. Further analysis showed that local differences were primarily influenced by the home geometry rather than the specific behavioral properties typically associated with a home cage. Specifically, the introduction of a plain box with similar geometrical properties to the home cage resulted in very similar changes in grid fields, even though the elicited behaviors were quite different. The model predicts the same behavior because it assumes that the effect of the home cage on predictive maps is primarily due to the way it alters transitions in relation to an open field, and because the DR (unlike the SR) codes predictive relationships in a way that is invariant to the expressed behavioral policy.

**Figure 7.**
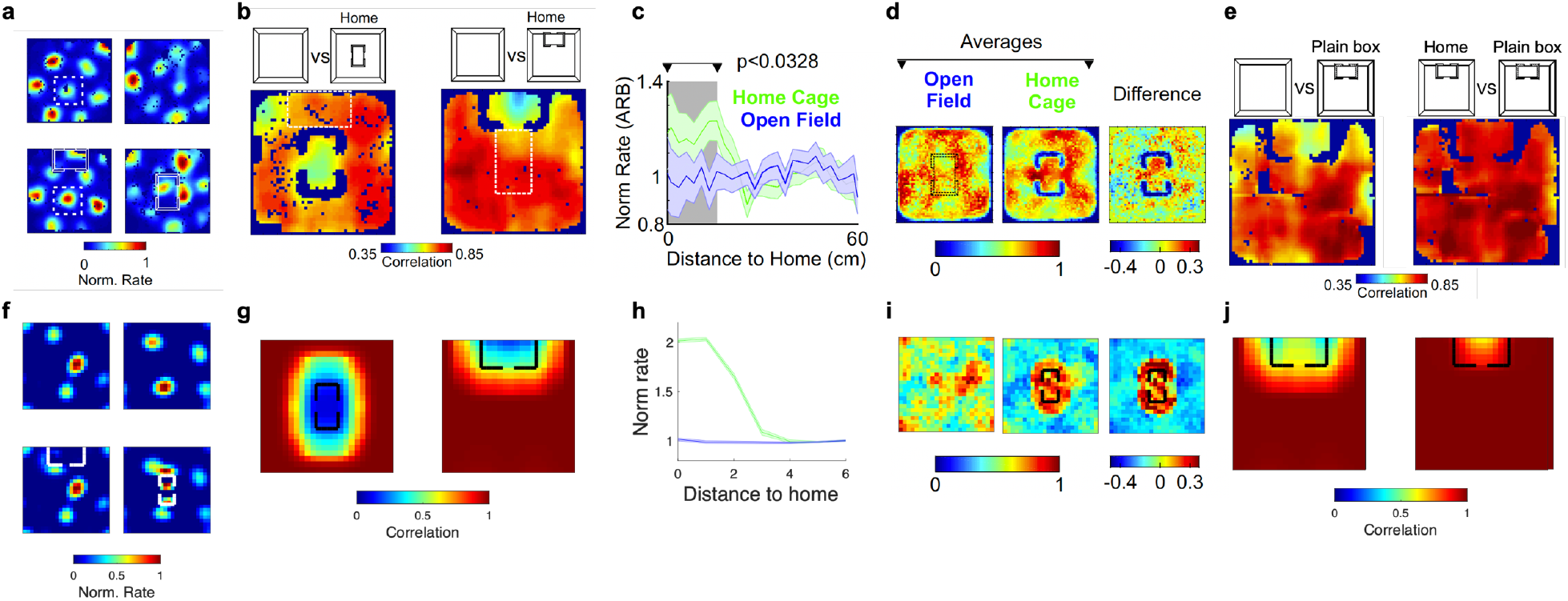
Grid maps change locally near the home cage. **a**. In the experiment by Sanguinetti-Scheck and Brecht, grid cells are recorded before and after introduction of a home cage. Here, two example normalized grid fields are plotted for two different home cage locations (north or middle). **b**. Sliding window spatial correlation map averaged across all cells is plotted for two different locations of the home cage. The impact of introduction the home cage is mostly limited to locations near the cage. **c**. The peak normalized firing map for all cells is plotted as a function the distance between each cell’s peak location and the center of the home cage. The local impact of the home cage on grid maps is also evident in the peak normalized rate map. **d**. The spatial average of the peak normalized rates is illustrated for both the open field and the home center condition, with the difference also displayed. The same effect is apparent in this comparison. **e**. Sliding window spatial correlation map averaged across all cells is plotted for another experimental condition in which a plain box is introduced with the same size as the home cage is introduced. The effect of the plain box is very similar to that the home cage, revealing that changes in grid cells are driven by the geometry rather than the home valence. **f-j**. Simulation of experimental results in **a-e** by the model. The simulation results in **f-j** were repeated 50 times with different randomization seeds and the average has been plotted. Errorbars in **h** indicate standard error across simulations. Data in **a-e** is published by Sanguinetti-Scheck and Brecht.

Of course, in environments where barriers alter almost the entire space relative to open areas, we expect more global remapping. For example, Derdikman and colleagues recorded grid cells in a hairpin maze environment, and the grid map was substantially distorted. Notably, a significant number of grid cells showed opposite patterns for alternating arms (Derdikman et al., 2009) suggesting a repetitive, compositional construction. The model predicts the same behavior, as the extent to which the barriers changed the open space is quite extensive here (Figure 8), and reflects the repetitive, compositional structure of the perturbation. Moreover, Derdikman et al. found that grid cells “reset” at turning points, a finding that is not consistent with classical global space coding theory of grid cells, but it is consistent with the theory that grid cells encode a cognitive map that predicts “fragmentation” in the predictive map. Finally, as we pointed out in our previous study (Piray & Daw, 2021), unlike the eigenvectors of the SR, experimental data support an implementation of predictive maps that is influenced by actual physical barriers but, in contrast, is relatively stable with respective to stereotypic behavioral policies. For instance, Derdikman and colleagues also tested whether grid cells are affected when rats trained to run equivalent hairpin pattern without any actual barriers (Derdikman et al., 2009). They found that grid fields in this case are more similar to the open field grids than the ones observed in the actual hairpin maze environment, a finding that is well explained by our model (again, due to the off-policy character of the DR), but not by policy-dependent predictive map models (Stachenfeld et al., 2017).

**Figure 8.**
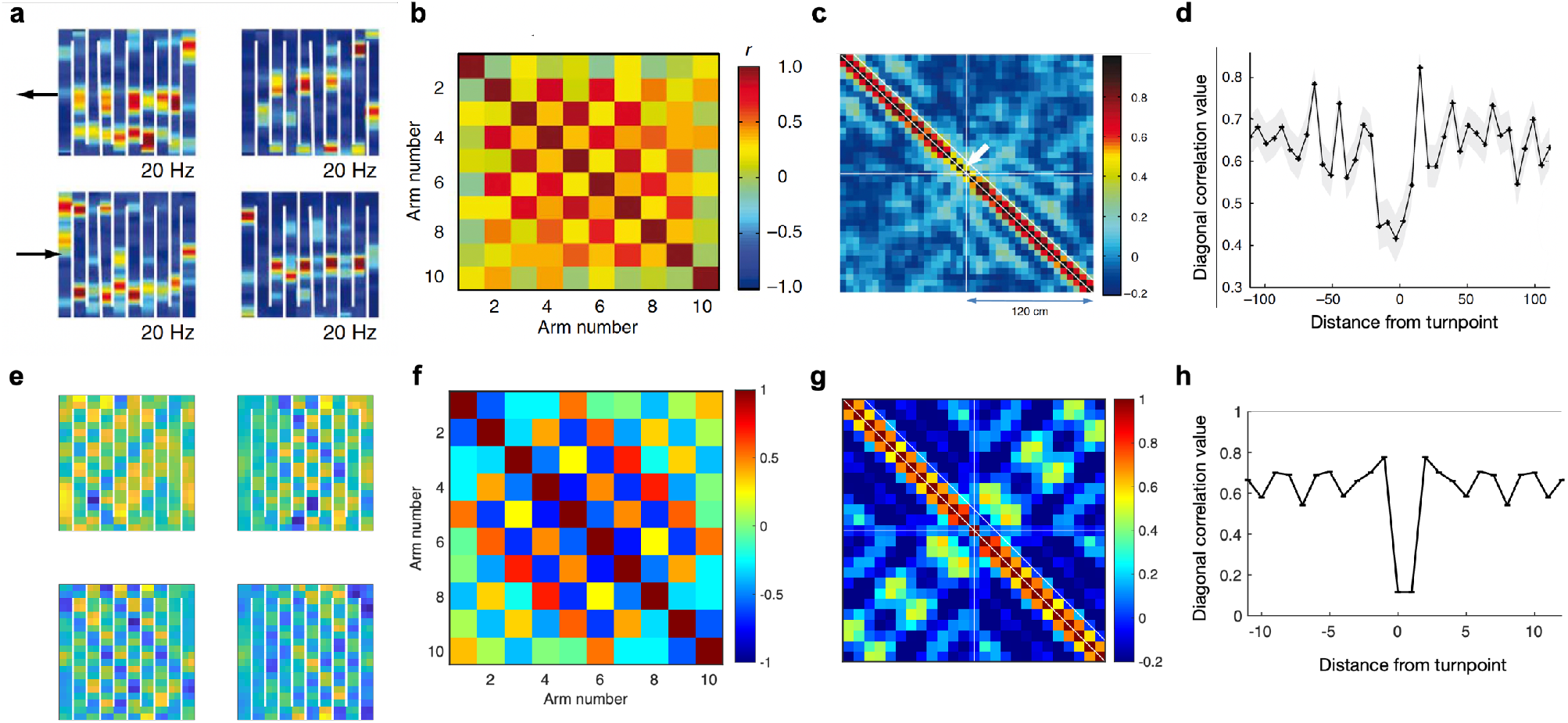
Predictive representations dramatically remap if extensive barrier is introduced in the environment. **a**. Firing rate of two example grid cells recorded in the hairpin-like environment show alternating firing pattern between different arms. **b**. Correlation between different arms across all recorded grid cells. **c**. Spatial correlation map is plotted, which is calculated based on population vectors of firing rates at the turning point and in 6-cm bins on both sides of the turning point. Directionally tuned grid cells were excluded from the population vectors. **d**. Correlation derived from the adjacent position bins selected from the diagonal section labeled in **a** is plotted. **e-h**. Simulation of experimental results in **a-d** by the model. The simulation results in **e-h** were repeated 50 times with different randomization seeds and the average has been plotted. Errorbars in **h** indicate standard error across simulations. Data in **a-d** is published by Derdikman and colleagues.

A final point about grid cells is their sensitivity to changing goals. Although it had been thought that grid fields are stable relative to the animal’s goal, recent studies suggest that goals subtly modulate grid fields (Boccara et al., 2019; Butler et al., 2019). It is not clear how this can be explained by previous models, even by predictive accounts, particularly the nuanced and subtle changes observed in grid maps. This is because the SR and its eigenvectors tend to change dramatically when a new goal is introduced, as this fundamentally alters the policy. Interestingly, though, goals in the current model do have a special status, which is similar to barriers, because the model concerns the so-called “episodic” RL setting, in which episodes terminate (and predictions are truncated; the next episode begins), whenever a goal is reached. Thus, our model distinguishes “terminal” states (containing goals) from other states, and if a state becomes a goal (terminal), this changes the transition probability between the goal state and its neighbors in an open field. Intuitively speaking, the goal state is similar to a one-way barrier because as soon as the agent gets to the goal, it does not move back as the planning episode is completed. Thus, we can similarly formalize the effect of the “goal component” on the predictive map. This aspect of the model accounts for recent studies that investigated the role of goals on grid fields (Boccara et al., 2019; Butler et al., 2019; Rueckemann et al., 2021). For example, Boccara and colleagues tested how grid fields change when rats are trained to daily learn three new reward locations on a cheeseboard maze (Boccara et al., 2019). They found a local remapping in grid cells after learning, in which grid fields mainly changed near the goal locations, with fields getting closer to the goal location. This also led to a reduction in the hexagonal grid pattern, consistent with a distortion of grid fields. The model shows the same behavior (Figure 9).

**Figure 9.**
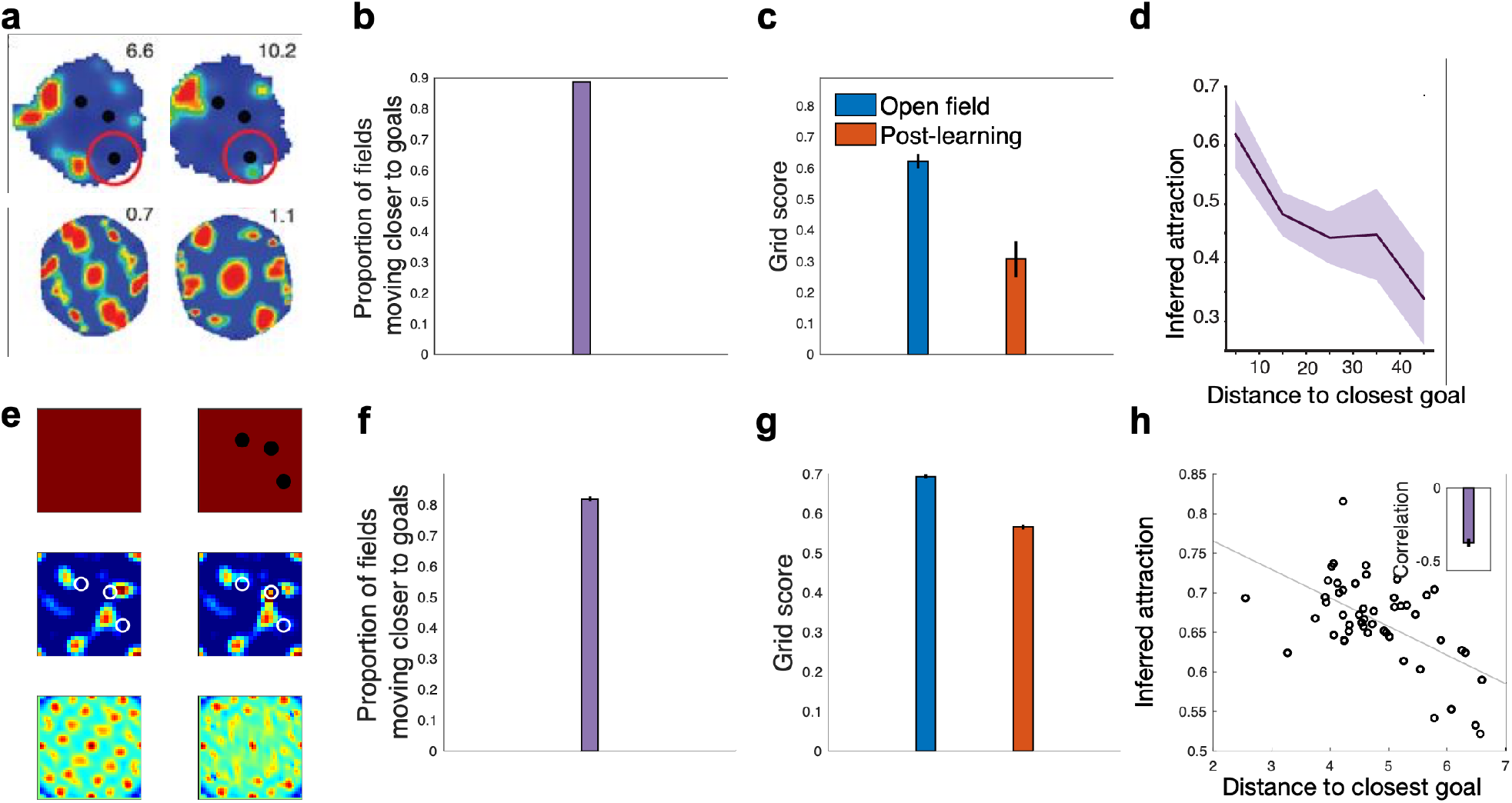
Introduction of goals locally modulate grid fields. **a**. Firing rate of an example grid cell is shown before and after learning the goal locations (blacks dots). Introduction of goals led to grid cells moving closer to goals. **c**. Grid score for the open field and after learning the goal locations is plotted. **d**. The degree of attraction of grid fields is inversely related to the distance of their peak from the closest goal. Simulation of experimental results in **a-d** by the model. The simulation results in **e-h** were repeated 50 times with different randomization seeds and the average has been plotted. Errorbars indicate standard error across simulations. Data in **a-d** is published by Boccara and colleagues.

## Discussion

Compositionality is an important aspect of a flexible system, whether biological or artificial. Compositionally thus seems particularly relevant to the concept of cognitive map, which is supposed to organize knowledge in a way that is generalizable across tasks and environments and enables flexible behavior, such as planning routes or taking novel shortcuts (Tolman, 1948). Neurons within MEC, such as grid cells and object vector cells, have been proposed as the neural substrates of the cognitive map (Behrens et al., 2018; Bellmund et al., 2018; McNaughton et al., 2006; Schacter et al., 2007), and have a number of properties that seem to suggest they represent different separable components of such a map. However, it has remained elusive how such a cognitive map could support flexible planning while being created compositionally. This includes both traditional spatial metric theories of MEC grid cells (Burak & Fiete, 2009), as well as newer models in which MEC represents an abstract state space (Behrens et al., 2018; Sorscher et al., 2023; Whittington et al., 2020) or a predictive SR (Piray & Daw, 2021; Stachenfeld et al., 2017). While some of these models emphasize one or the other of these features, none of them reconciles the cognitive map’s utility for planning with the need for it to be built compositionally from task elements, such as objects, barriers, and goals. Here, we have proposed a predictive model for planning that is also created compositionally by assembling representations related to objects and goals. This also gives a clear and explicit computational interpretation to the properties of different spatially tuned neurons in MEC.

The new model is based on our linear RL theory, which posits that, given a long-range predictive map encoding closeness between all states under a random policy, optimal planning can be accurately and efficiently approximated for any goal (Piray & Daw, 2021). Moreover, our approach leverages four key computational features: 1) Objects, barriers, and goals can be viewed as perturbations disrupting the relational structure of open space. 2) Their effects on long-range predictive maps can be efficiently computed using translation- and rotation-invariant matrices. These are efficient as their size depends on the object, not full state space. 3) Long-range maps can be efficiently constructed by combining these compact predictive object representations. 4) There exists an efficient set of basis functions for representing the cognitive map by combining PORs with the eigenvectors of the open space.

Our computational approach lays a substantially improved foundation for understanding the function of different cells within the MEC, including object vector cells or border cells encoding elements of the task space, as well as grid cells providing basis function of the compositional predictive map. In particular, we showed that the proposed code for representing objects resembles remarkable similarity to object vector cells, an important class of cells discovered recently in the MEC (Høydal et al., 2019). Similar to object vector cells, PORs are translation- and rotation-invariant, and their field size depends on the object size. The model further provides a computational mechanism of how grid cells might reflect basis functions for cognitive maps. In fact, by combing PORs with eigenvectors of the open space, one can obtain a low-dimensional representation of the proposed compositional predictive maps, a process that can be done efficiently using a simple neural network with known weights. This makes a critical prediction, particularly about the effects of barriers and goals on grid fields. Specifically, we anticipate that effects of barriers and goals on grid fields remain relatively local, a prediction that is well supported by empirical data (Ginosar et al., 2023).

Our theory relates to earlier work by Stachenfeld et al. proposing that grid cells encode eigenvectors of the SR (Stachenfeld et al., 2017). The SR constitutes a predictive map of task space and can be seen as a policy-dependent equivalent to the DR utilized here (Dayan, 1993; Russek et al., 2017). In many ways, our work builds upon this idea. However, the Stachenfeld et al. model faces major computational and empirical challenges. Computationally, it is unclear how such a code can be implemented in an efficient way given that all eigenvectors must be recomputed with any change to goals or environment structure. Notably, the SR’s size is quadratic in the number of states in the environment – thus computing its eigenvectors even once is often intractable in a large state space, let alone repeated full computation of eigenvectors with every change. The SR also makes predictions about the neural code that are contradicted by data, in that eigenvectors of the SR would exhibit extensive remapping when barriers, objects, or goals are introduced. This contrasts with experimental data showing local grid remapping in such situations (Ginosar et al., 2023), especially in naturalistic environments (Boccara et al., 2019; Butler et al., 2019; Ginosar et al., 2021; Krupic et al., 2018; Sanguinetti-Scheck & Brecht, 2020).

Our new theory also relates to a recent but growing body of work proposing probabilistic, graph-like computational models of MEC cognitive maps, which argue that MEC encodes relational structure between different states in the world (Behrens et al., 2018; Peer et al., 2021; Whittington et al., 2020). Our model extends this line of work in two critical ways. First, these previous models are often agnostic to planning, because they do not explicitly model reward as an entity organisms seek to maximize. Indeed, because they generally represent the one-step map, building planning over this representation is computationally quite laborious. Our model creates cognitive maps that, together with the linear RL framework, provide an efficient method for planning. Second, while one of these models (REF TEM) does capture a sort of compositionality, in that it based on learning to generalize relational graph connectivity motifs across different environments, this property emerges only implicitly and somewhat opaquely by training a black-box neural network via backpropagation. This in turn limits the functional interpretation of units within the simulated networks, which are observed to resemble those in MEC. Our current model additionally provides an explicit, algebraically derived method for constructing maps compositionally from individual components representing barriers and goals. Thus, our model builds on these graph-based cognitive mapping theories, while more explicitly exposing the map’s compositional structure, its role in planning, and the computational role of different neural responses.

The current theory expands on our prior linear RL model for planning, which efficiently reduces planning to constructing predictive maps (Piray & Daw, 2021). One key feature of the model, which distinguishes it from similar predictive algorithms like the SR, is that it is policy independent. That is, it represents predictions about long-run state encounters under a fixed, “default” policy (e.g., a random walk), but it can be used to solve approximately for the value function under any goal-optimized policy. In the earlier work we showed how this enabled various types of transfer learning relevant to cognitive maps; in the current work it is crucial computationally for enabling the stable compositional construction of the map and empirically for explaining various phenomena related to the stability of grid fields. Overall, the current theory complements our linear RL model by delineating how these maps can be created compositionally from task elements, such as barriers, objects, and goals. Together, this provides an efficient, comprehensive, and psychobiologically plausible theory of model-based planning.

When facing two or more objects, we demonstrate that a first-order approximation of the solution can be obtained through compositional combination of the representations related to each individual object. We further propose a learning algorithm for modeling interactions between objects. Importantly, this algorithm is efficient as its starting point is already a decent approximation of the full solution, as depicted in Fig 1. Additionally, it is computationally efficient since the object-related representation matrices scale only with object size rather than environment size. This contrasts with much of the previous work on SR, in which learning the map at least required updating entire rows or columns of the SR matrix, which is the size of the state space (Russek et al., 2017). Thus, by leveraging compositionality and initially approximating multi-object solutions from single object representations, our approach provides an efficient framework for learning object interactions and constructing multi-object cognitive maps. Although the neural basis of this process remains unclear, mental simulation through hippocampal replay may offer insights into this process. During periods of wakefulness, place cells exhibit rapid, sequenced reactivations known as awake replays (Davidson et al., 2009; Diba & Buzsáki, 2007; Foster & Wilson, 2006; Pfeiffer & Foster, 2013), which have been linked to planning (Jadhav et al., 2012; Pfeiffer & Foster, 2013). Initial evidence indicates these hippocampal replays remain largely stable even when barriers and goals change in an environment (Widloski & Foster, 2022). This suggests replays could provide a neural mechanism for learning compositional cognitive maps in a flexible manner. Though the neural mechanisms are still unknown, awake replays represent a promising phenomenon to explore in future research on the neuroscience of compositional mapping.

The current theory provides a comprehensive and biologically realistic computational framework for constructing cognitive maps useful for model-based planning in the brain. This establishes a foundation for future work investigating the neural substrates underlying cognitive mapping. Although we have focused on the MEC’s role in cognitive maps, map making likely involves multiple neural systems rather than a single localized mechanism. Different systems may construct maps in distinct contexts, like social, emotional, or spatial. While our framework models interactions between subprocesses of map making, we do not imply these computations are localized to specific regions. More realistically, various neural systems participate in different computational subprocesses of cognitive mapping. Overall, our theory provides a unifying computational account of cognitive mapping and model-based planning while remaining compatible with distributed, systems-level neural implementation. This integrative approach opens promising directions for examining the neural mechanisms of how diverse brain areas cooperate to enable efficient map making, and flexible planning.

This work also aligns with emerging theoretical and empirical research tapping into neural basis of representation learning, or how human and other animals construct task representations that allow efficient decision making (Niv, 2019). In particular, our model suggests that the way that an object or barrier changes the relational structure of the state space is of primary importance for planning, whereas the sensory features of such objects are irrelevant and ignored by our model’s representations. This is not only compatible with representations of object vector cells in the MEC as shown here, but it is also consistent with recent evidence suggesting that neural representations in MEC and medial prefrontal cortex are relatively invariant to sensory aspects of task elements (Park et al., 2021; Whittington et al., 2020). While testing representational models remains challenging, recent advances demonstrate feasibility. For instance, combining neuroimaging and representational similarity analysis techniques (Kriegeskorte et al., 2008; Nili et al., 2014) has elucidated how the hippocampus and orbitofrontal cortex encode social cognitive maps (Park et al., 2020, 2021). Further developing such techniques will be key to unlocking the computational principles represented in neural systems. An integrative interplay between computational modeling and representation-focused experimentation will likely offer new insights into the algorithms underlying cognitive map making and model-based planning.

Within our model, maps are created by integrating object related representations with the basis functions of the open space. This suggests local interactions between object vector cells and grid cells. More specifically, our approach suggests that the grid cells might receive inputs from other cell types, such as object vector cells (Moser et al., 2015). Although the neural mechanisms underlying the implementation of the proposed grid code in the MEC are still unclear, recent advances have brought us to the point where these underlying mechanisms can soon be elucidated (Ginosar et al., 2023; Tukker et al., 2022). In particular, new electrophysiological techniques now enable simultaneous recording of hundreds of MEC neurons during navigation (Gardner et al., 2022). It would be important to leverage these techniques to conduct multi-neuronal recordings of different MEC cell types as animals move through environments containing objects and goals. Analyzing the interactions between cell types under these conditions could reveal how the grid code emerges from local microcircuit computations and whether those are consistent with the quantities predicted by the model.

## Methods

### Linear RL

We start with a review of the linear RL model (Piray & Daw, 2021), which considers finite-horizon Markov decision problems with deterministic dynamics, such as mazes. The linear RL model maximizes long-term gain defined as a linear function of the reward for each state. In particular, if *r*(*s*) is the reward in state *s*, and *λ* > 0 is a constant called control cost, then gain, *g*, is defined as *g*(*s*) = *r*(*s*) − *λ*KL(*π*||*π*^*d*^). Here, KL(*π*||*π*^*d*^) is the Kullback-Leibler divergence between the decision policy, *π*, and a default policy *π*^*d*^ (the policy under which the map is made, e.g. a random walk). This is a measure of how two probability distributions diverge from each other. It is zero if *π* equals *π*^*d*^, and otherwise, it is positive. Throughout this work, we always assume a uniform default policy, which is equivalent to moving to all successor states with equal probability.

It is then straightforward to demonstrate that the optimal value function for this problem, *v*^*^(*s*), is analytically solvable. (It can also be viewed as an approximation to value of the original RL problem, without control costs.) We start by defining the one-step state transition matrix, denoted as **T**. The (*i, j*) element of **T** represents the probability of transitioning from state *i* to state *j* under the default policy (i.e. probability of the action under the default policy that leads to transition from state *i* to state *j*). We further distinguish states into terminal states (containing goals) and all other, nonterminal states. Next, we define **t** = **T**_*NT*_ exp(*r*_*T*_/*λ*), a vector that depends only on the transition to the terminal state, **T**_*NT*_ and its reward, *r*_*T*_. Now, if **v**^*^ is the vector of optimal values across all nonterminal states, then exp(**v**^*^/*λ*) = **D**_*NN*_**t**. Here, **D**_*NN*_ is a subblock of the DR matrix, **D**, corresponding to nonterminal states. The DR is defined as:

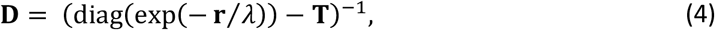

where **r** is the reward vector across all states (assuming, without loss of generality, that reward at terminal states is not 0). Thus, the DR, **D**, is a matrix where the (*i, j*) element indicates the expected occupancy of state *j*, discounted by exponential state rewards (which is negative), when starting from state *i* and following the default policy. It differs from the SR, which represents expected occupancy under the decision policy (and thus changes whenever the optimal values change, such as when goals move). The DR matrix **D** is the main focus of the current work.

### Compositional map making

Suppose that **D**_*os*_ is the DR of the open space for a random walk (if one takes actions randomly according to a uniform transition probability.) In practice, it can be efficiently defined by calculating the DR for a significantly larger environment and then extracting the DR corresponding to a subregion from the middle. The subregion should be of a size equivalent to the task environment. We assume that all state costs are equally given by the constant *c*. Thus, if **T**_*os*_ is the transition matrix under the random walk, **I** is the identity matrix with size equal to **T**_*os*_, and **L**_*os*_ = exp(*c*) **I** − **T**_*os*_, then 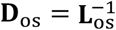.

Now consider an object *o* that changes the transition matrix to **T**_*o*_. It is easy to see that **T**_*o*_ can be written as a function of **T**_*os*_, and two low-rank matrices:

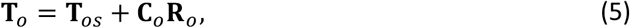

where **C**_*o*_ and **R**_*o*_ are, respectively, *S* × *O* and *O* × *S* matrices, where *O* is the number of states whose transition changed by introduction of the object, and *S* ≫ *O* is the number of states in the open space environment. Since the object only changes the transition for states that are neighbor to one of the states that is blocked by the object, *O* is essentially close to the size of object. Both **C**_*o*_ and **R**_*o*_ are sparse matrices with nonzero elements only on rows or columns associated with the object and its neighbor states. In particular, if {*s*_1_, *s*_2_, …, *s*_*O*_} represents the set of states whose transitions were affected by the introduction of the object, the *j*th column of matrix **C**_*o*_ is a binary vector with its *j*th element set to one. Additionally, the *j*th row of matrix **R**_*o*_ is defined as the *j*th row of matrix **T**_*o*_ − **T**_*os*_.

Now, if **L**_*o*_ = exp(*c*) **I** − **T**_*o*_ = **L**_*os*_ − **C**_*o*_**R**_*o*_, we can use Woodbury matrix inversion identity (Hager, 1989) to write 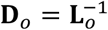 as a function of **D**_*os*_ :

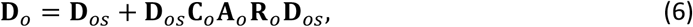

where **A**_*o*_ is a *O* × *O* matrix representing the POR for the object *o*. If we define **Z**_*o*_ = **R**_*o*_**D**_*os*_**C**_*o*_, then **A**_*o*_ is given by:

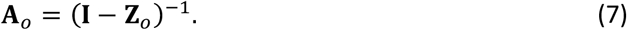

If |**Z**_*o*_| < 1, **A**_*o*_ is given by Neumann series:

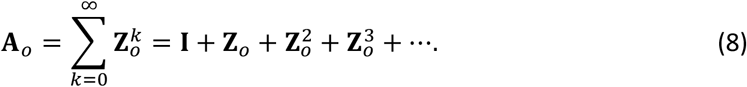

Thus, **A**_*o*_ ≃ **I** + **Z**_*o*_ provides a first-order approximation of **A**_*o*_.

Now, let’s consider an environment that includes two distinct objects, labeled 1 and 2. We assume these objects are distinguished in such a way that there is no single state whose transition is affected by both objects. As a result, the lookup table matrices for the composition of the objects can be expressed as column- and row-concatenation of their associated matrices, i.e., [**C**_1_ **C**_2_] and 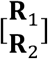.

By combining Equations 6 and 8, it is easy to see that Equation 2 provides an approximation for DR matrix **D** of the new environment. This approximation approaches the exact solution as the distance between the two objects increases.

### Learning PORs

Equation 2 provides a first-order approximation of the POR for two (or more) objects. Building on this, we have developed an algorithm for learning the POR that is inspired by temporal difference learning rules from RL. First, we define matrix **Z** based on the lookup table matrices, **C** and **R**, for the composition of the two objects. These are defined as column- and row-concatenation of their corresponding matrices, i.e., **C** = [**C**_1_ **C**_2_] and 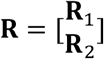. Thus, we define **Z** = **CD**_*os*_**R**. If **A** is initially set to **I** + **Z**, and 0 < *α* < 1 is a scalar parameter representing the step-size, we iteratively update our POR estimate, **A**, using the following equation:

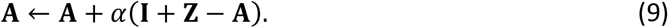

It can be demonstrated that **A** consistently converges to the exact solution, i.e., (**I** − **Z**)^−1^, with each update, as long as the Neumann series (Equation 8) exists for **Z**, i.e., |**Z**| < 1. In addition, this algorithm is highly efficient for two reasons. Firstly, the initial point provides a strong approximation of the POR. Secondly, unlike classical temporal difference algorithms for updating the SR (Dayan, 1993; Russek et al., 2017), this one operates within the object space, given that the sizes of matrices **A** and **Z** are object dependent.

### Simulation of object vector cells

Our model characterizes object vector cells as encoding the projected POR, which captures the change in the DR induced by the introduction of an object into the environment (Equation 1), i.e., **D**_*o*_ = **D**_*os*_**C**_*o*_**A**_*o*_**R**_*o*_**D**_*os*_ Specifically, we model object vector cells as a row vector of the POR corresponding to the specific state, let’s say *s*, associated with the particular distance and direction from the object to which the cell is tuned (due to the vector coding (Bicanski & Burgess, 2020)). Thus, the object vector cell code is given by the *s*th row of the **D**_*o*_. The environment for simulating the object vector cells was created by discretizing the plane into a network of triangular lattice points. It was assumed that every state is connected to any other state in its vicinity determined by parameter *r*. This is equivalent to the assumption that the width place code is given by *r*. Additionally, we assumed that a barrier representation for object vector cells requires the object to be visible from the agent’s position, aligning with empirical observations (Høydal et al., 2019). This was modeled by assuming that a transition was possible from the agent’s position to the object’s location before the introduction of the object, and the object subsequently obstructed that transition.

### Simulation of grid cells

The open space grid map was modeled using the framework and code developed by Dordek et al. (Dordek et al., 2016). The open space environment consisted of uniformly distributed 2D Gaussian-shaped place cells, organized in a grid pattern. The activity of each place cell *i* at location **x** was modeled as a normal distribution with mean *c*_2_ and variance 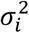. The mean indicates the center of the place field for cell *i*, while the variance 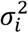 determines the width of the place field. The place field centers were uniformly distributed across the environment, and all had the same width. The activity of *N* place cells over *T* time-points was governed by a random uniform policy dictating the agent’s movement. This produced a *N* × *T* matrix of place cell activities. The eigenvectors were computed by running an eigen-decomposition (or, equivalently, a principal component analysis) on this matrix. As in Dordek et al. (Dordek et al., 2016), a nonnegativity constraint was applied. The first 50 principal components calculated using the Oja rule (Oja, 1982) were used in our simulations. Since these components are calculated for the covariance matrix of place cell activities, we used the covariance matrix as **D**_*os*_ to calculate **G** in Equation 3.

The task environment for each grid field simulation was created as a 2D square grid representing the tabular state space. For each task, barriers were modeled by blocking the corresponding transitions between states.

### Simulation details

For the model simulation in Figure 1, we assumed a 20×20 square maze environment. The cost for all states was set to 0.1, with the goal state having 0 cost. To minimize the effects of the borders on the open space DR, we first computed the DR for a significantly larger open space (100×100) and subsequently selected the subsection that corresponds to our maze environment. The step-size in Equation 10 was 0.3. For implementing the SR, we assumed the reward vector was given (similar to the assumption for our model), and the algorithm only learned the SR map using a step size of 0.2. Actions were selected based on a softmax policy with decision noise of 1. Since this was a finite horizon problem (i.e. planning towards a goal state), the discount factor was set to 1. For all simulations related to object vector cells presented in Figure 2-4, the environment size was 20, and *r* = 2 was used as the place code width. The cost for all states was 0.1.

For grid field simulations, we generally used the same set of parameters as the open space grid field. For the simulation in Figure 6, each compartment was 30×30, and *σ* was 0.35. To calculate the sliding spatial correlation, we assumed a box size width of 6, ensuring that its ratio compared to the maze size roughly resembles that of the original experiment. For the simulation presented in Figure 7, the environment was 25×25 and *σ* was 0.35. It was assumed that states at the center of the home cage serve as terminal states, an assumption not made for the plain box. The box size for calculating the spatial correlation was 3. The normalized firing rate was computed as the average of fields within the corresponding distance surpassing a threshold, divided by the same metric across the entire maze in the open space environment. The threshold was defined as the average of all grid vectors in the open space. The grid vectors plotted in Figure 7F were normalized by their maximum values, resulting in a range of values from 0 to 1. For calculating all spatial correlation maps, all states at the edge of the environment were excluded.

For the hairpin experiment presented in Figure 8, the environment was 20×20 and *σ* was 0.35. To calculate the correlation matrix between hairpin arms, the median correlation across all grid vectors exhibiting a significant alternating pattern (based on Wilcoxon rank difference) was computed for each arm. The turning point activity analysis involved computing the population activity of firing rates within paths containing a turning point and adjacent bins on both sides. Following the original study, our focus was on the activity of grid vectors that did not exhibit alternating patterns when calculating the turning matrix activity. As a result, the population firing rate was defined as the average of all such grid vectors. In our simulation, the turning point actually encompassed four states: one on one side of the barrier, two situated just next to the barrier but unblocked, and one on the other side of the barrier. The population activity at the turning point was determined as the median activity across all four states associated with the turning point. Furthermore, population activity across 12 states along each side of the barrier was also considered, resulting in vectors with a length of 25. The resulting population activity matrix was organized as a 25×9 matrix, where 9 represents the number of turning points in the environment. The turning correlation matrix was then calculated by computing correlations across columns of this matrix, resulting in a 25×25 matrix. This analysis was repeated 50 times with different randomization seeds, and the mean matrices were plotted in Figure 8f-h.

For the simulation presented in Figure 9, the environment was 25×25 and *σ* was 0.35. For calculating the number of grid fields moving closer to the goals, we first calculated the locations around the goal (all states within the radius of 2.5 of the center of a goal location). The peak activity near goals per grid vector was defined as the mean of all grid fields whose value is larger than a threshold. The threshold here was defined based on the mean normalized activity across all grid vectors, where the normalized activity is the median of the grid vector divided by its maximum. Mean number of vectors whose peak activity near goals was larger after the introduction of goal was the index plotted in Figure 9f. The grid score (Sargolini et al., 2006) was calculated using the code implemented by Dordek et al. (Dordek et al., 2016). The mean of grid scores was calculated across all vectors with an open space grid score eceeding 0.5 and used in Figure 9g. For analyzing attraction as a function of distance, we took into account all active fields, defined as those with normalized activity (ranging between 0 and 1) exceeding half. The mean distance between the field peak location and the center of each of the three goals served as the corresponding distance metric for that active field. For each active field, attraction vector was defined if the median activity of the vector in the vicinity of the peak location increased following the introduction of the goals, considering any location within a distance of less than 2.5 as part of the field’s vicinity. Field attraction plotted in Figure 9h was defined as the mean of attraction vector. This analysis was repeated 50 times with different randomization seeds, and the mean metrics were plotted in Figure 9f-h.

The control cost parameter, *λ*, in Equation 4 was set to 1 for all simulations.

## References

Andersson, S. O., Moser, E. I., & Moser, M.-B. (2021). Visual stimulus features that elicit activity in object-vector cells. Communications Biology, 4(1), 1219. 10.1038/s42003-021-02727-5

Behrens, T. E. J., Muller, T. H., Whittington, J. C. R., Mark, S., Baram, A. B., Stachenfeld, K. L., & Kurth-Nelson, Z. (2018). What Is a Cognitive Map? Organizing Knowledge for Flexible Behavior. Neuron, 100(2), 490–509. 10.1016/j.neuron.2018.10.002

Bellmund, J. L. S., Gärdenfors, P., Moser, E. I., & Doeller, C. F. (2018). Navigating cognition: Spatial codes for human thinking. Science (New York, N.Y.), 362(6415), eaat6766. 10.1126/science.aat6766

Bicanski, A., & Burgess, N. (2020). Neuronal vector coding in spatial cognition. Nature Reviews. Neuroscience, 21(9), 453–470. 10.1038/s41583-020-0336-9

Bienenstock, E., Geman, S., & Potter, D. (1996). Compositionality, MDL priors, and object recognition. Proceedings of the 9th International Conference on Neural Information Processing Systems, 838–844.

Boccara, C. N., Nardin, M., Stella, F., O’Neill, J., & Csicsvari, J. (2019). The entorhinal cognitive map is attracted to goals. Science (New York, N.Y.), 363(6434), 1443–1447. 10.1126/science.aav4837

Burak, Y., & Fiete, I. R. (2009). Accurate Path Integration in Continuous Attractor Network Models of Grid Cells. PLOS Computational Biology, 5(2), e1000291. 10.1371/journal.pcbi.1000291

Butler, W. N., Hardcastle, K., & Giocomo, L. M. (2019). Remembered reward locations restructure entorhinal spatial maps. Science (New York, N.Y.), 363(6434), 1447–1452. 10.1126/science.aav5297

Chang, M. B., Ullman, T., Torralba, A., & Tenenbaum, J. B. (2017). A Compositional Object-Based Approach to Learning Physical Dynamics (1612.00341). arXiv. 10.48550/arXiv.1612.00341

Davidson, T. J., Kloosterman, F., & Wilson, M. A. (2009). Hippocampal replay of extended experience. Neuron, 63(4), 497–507. 10.1016/j.neuron.2009.07.027

Daw, N. D., Niv, Y., & Dayan, P. (2005). Uncertainty-based competition between prefrontal and dorsolateral striatal systems for behavioral control. Nature Neuroscience, 8(12), 1704–1711. 10.1038/nn1560

Dayan, P. (1993). Improving Generalization for Temporal Difference Learning: The Successor Representation. Neural Computation, 5(4), 613–624. 10.1162/neco.1993.5.4.613

de Cothi, W., & Barry, C. (2020). Neurobiological successor features for spatial navigation. Hippocampus, 30(12), 1347–1355. 10.1002/hipo.23246

Derdikman, D., Whitlock, J. R., Tsao, A., Fyhn, M., Hafting, T., Moser, M.-B., & Moser, E. I. (2009). Fragmentation of grid cell maps in a multicompartment environment. Nature Neuroscience, 12(10), 1325–1332. 10.1038/nn.2396

Deshmukh, S. S., & Knierim, J. J. (2013). Influence of local objects on hippocampal representations: Landmark vectors and memory. Hippocampus, 23(4), 253–267. 10.1002/hipo.22101

Diba, K., & Buzsáki, G. (2007). Forward and reverse hippocampal place-cell sequences during ripples. Nature Neuroscience, 10(10), Article 10. 10.1038/nn1961

Dordek, Y., Soudry, D., Meir, R., & Derdikman, D. (2016). Extracting grid cell characteristics from place cell inputs using non-negative principal component analysis. eLife, 5, e10094. 10.7554/eLife.10094

Fodor, J. A., & Pylyshyn, Z. W. (1988). Connectionism and cognitive architecture: A critical analysis. Cognition, 28(1–2), 3–71. 10.1016/0010-0277(88)90031-5

Foster, D. J., & Wilson, M. A. (2006). Reverse replay of behavioural sequences in hippocampal place cells during the awake state. Nature, 440(7084), 680–683. 10.1038/nature04587

Frankland, S. M., & Greene, J. D. (2020). Concepts and Compositionality: In Search of the Brain’s Language of Thought. Annual Review of Psychology, 71, 273–303. 10.1146/annurev-psych-122216-011829

Gardner, R. J., Hermansen, E., Pachitariu, M., Burak, Y., Baas, N. A., Dunn, B. A., Moser, M.-B., & Moser, E. I. (2022). Toroidal topology of population activity in grid cells. Nature, 602(7895), Article 7895. 10.1038/s41586-021-04268-7

George, D., Rikhye, R. V., Gothoskar, N., Guntupalli, J. S., Dedieu, A., & Lázaro-Gredilla, M. (2021). Clone-structured graph representations enable flexible learning and vicarious evaluation of cognitive maps. Nature Communications, 12(1), Article 1. 10.1038/s41467-021-22559-5

Gershman, S. J. (2018). The Successor Representation: Its Computational Logic and Neural Substrates. The Journal of Neuroscience, 38(33), 7193–7200. 10.1523/JNEUROSCI.0151-18.2018

Gershman, S. J., Horvitz, E. J., & Tenenbaum, J. B. (2015). Computational rationality: A converging paradigm for intelligence in brains, minds, and machines. Science (New York, N.Y.), 349(6245), 273–278. 10.1126/science.aac6076

Ginosar, G., Aljadeff, J., Burak, Y., Sompolinsky, H., Las, L., & Ulanovsky, N. (2021). Locally ordered representation of 3D space in the entorhinal cortex. Nature, 596(7872), Article 7872. 10.1038/s41586-021-03783-x

Ginosar, G., Aljadeff, J., Las, L., Derdikman, D., & Ulanovsky, N. (2023). Are grid cells used for navigation? On local metrics, subjective spaces, and black holes. Neuron, 111(12), 1858–1875. 10.1016/j.neuron.2023.03.027

Glascher, J., Daw, N., Dayan, P., & O’Doherty, J. P. (2010). States versus rewards: Dissociable neural prediction error signals underlying model-based and model-free reinforcement learning. Neuron, 66(4), 585–595. 10.1016/j.neuron.2010.04.016

Goyal, A., & Bengio, Y. (2022). Inductive biases for deep learning of higher-level cognition. Proceedings of the Royal Society A: Mathematical, Physical and Engineering Sciences, 478(2266), 20210068. 10.1098/rspa.2021.0068

Greff, K., van Steenkiste, S., & Schmidhuber, J. (2020). On the Binding Problem in Artificial Neural Networks (2012.05208). arXiv. 10.48550/arXiv.2012.05208

Hager, W. W. (1989). Updating the Inverse of a Matrix. SIAM Review, 31(2), 221–239.

Hill, C. A., Suzuki, S., Polania, R., Moisa, M., O’Doherty, J. P., & Ruff, C. C. (2017). A causal account of the brain network computations underlying strategic social behavior. Nature Neuroscience, 20(8), 1142–1149. 10.1038/nn.4602

Ho, M. K., Abel, D., Correa, C. G., Littman, M. L., Cohen, J. D., & Griffiths, T. L. (2022). People construct simplified mental representations to plan. Nature, 606(7912), 129–136. 10.1038/s41586-022-04743-9

Høydal, Ø. A., Skytøen, E. R., Andersson, S. O., Moser, M.-B., & Moser, E. I. (2019). Object-vector coding in the medial entorhinal cortex. Nature, 568(7752), 400–404. 10.1038/s41586-019-1077-7

Hunt, L. T., Daw, N. D., Kaanders, P., MacIver, M. A., Mugan, U., Procyk, E., Redish, A. D., Russo, E., Scholl, J., Stachenfeld, K., Wilson, C. R. E., & Kolling, N. (2021). Formalizing planning and information search in naturalistic decision-making. Nature Neuroscience, 24(8), 1051–1064. 10.1038/s41593-021-00866-w

Jadhav, S. P., Kemere, C., German, P. W., & Frank, L. M. (2012). Awake Hippocampal Sharp-Wave Ripples Support Spatial Memory. Science, 336(6087), 1454–1458. 10.1126/science.1217230

Kinkhabwala, A. A., Gu, Y., Aronov, D., & Tank, D. W. (2020). Visual cue-related activity of cells in the medial entorhinal cortex during navigation in virtual reality. eLife, 9, e43140. 10.7554/eLife.43140

Kriegeskorte, N., Mur, M., & Bandettini, P. (2008). Representational Similarity Analysis – Connecting the Branches of Systems Neuroscience. Frontiers in Systems Neuroscience, 2, 4. 10.3389/neuro.06.004.2008

Krupic, J., Bauza, M., Burton, S., & O’Keefe, J. (2018). Local transformations of the hippocampal cognitive map. Science, 359(6380), 1143–1146. 10.1126/science.aao4960

Lake, B. M. (2019). Compositional generalization through meta sequence-to-sequence learning (1906.05381). arXiv. 10.48550/arXiv.1906.05381

Lake, B. M., & Baroni, M. (2023). Human-like systematic generalization through a meta-learning neural network. Nature, 623(7985), Article 7985. 10.1038/s41586-023-06668-3

Lake, B. M., Ullman, T. D., Tenenbaum, J. B., & Gershman, S. J. (2017). Building machines that learn and think like people. The Behavioral and Brain Sciences, 40, e253. 10.1017/S0140525X16001837

Liu, N., Li, S., Du, Y., Torralba, A., & Tenenbaum, J. B. (2023). Compositional Visual Generation with Composable Diffusion Models (2206.01714). arXiv. 10.48550/arXiv.2206.01714

McNaughton, B. L., Battaglia, F. P., Jensen, O., Moser, E. I., & Moser, M.-B. (2006). Path integration and the neural basis of the “cognitive map.” Nature Reviews Neuroscience, 7(8), Article 8. 10.1038/nrn1932

Meng, C., He, Y., Song, Y., Song, J., Wu, J., Zhu, J.-Y., & Ermon, S. (2022). SDEdit: Guided Image Synthesis and Editing with Stochastic Differential Equations (2108.01073). arXiv. 10.48550/arXiv.2108.01073

Momennejad, I., Russek, E. M., Cheong, J. H., Botvinick, M. M., Daw, N. D., & Gershman, S. J. (2017). The successor representation in human reinforcement learning. Nature Human Behaviour, 1(9), 680–692. 10.1038/s41562-017-0180-8

Moser, M.-B., Rowland, D. C., & Moser, E. I. (2015). Place Cells, Grid Cells, and Memory. Cold Spring Harbor Perspectives in Biology, 7(2), a021808. 10.1101/cshperspect.a021808

Nie, W., Vahdat, A., & Anandkumar, A. (2021). Controllable and Compositional Generation with Latent-Space Energy-Based Models (2110.10873). arXiv. 10.48550/arXiv.2110.10873

Nili, H., Wingfield, C., Walther, A., Su, L., Marslen-Wilson, W., & Kriegeskorte, N. (2014). A toolbox for representational similarity analysis. PLoS Computational Biology, 10(4), e1003553. 10.1371/journal.pcbi.1003553

Niv, Y. (2019). Learning task-state representations. Nature Neuroscience, 22(10), 1544–1553. 10.1038/s41593-019-0470-8

Oja, E. (1982). A simplified neuron model as a principal component analyzer. Journal of Mathematical Biology, 15(3), 267–273. 10.1007/BF00275687

O’Keefe, J., & Nadel, L. (1978). The Hippocampus as a Cognitive Map. Oxford University Press.

Park, S. A., Miller, D. S., & Boorman, E. D. (2021). Inferences on a multidimensional social hierarchy use a grid-like code. Nature Neuroscience, 24(9), 1292–1301. 10.1038/s41593-021-00916-3

Park, S. A., Miller, D. S., Nili, H., Ranganath, C., & Boorman, E. D. (2020). Map Making: Constructing, Combining, and Inferring on Abstract Cognitive Maps. Neuron, 107(6), 1226-1238.e8. 10.1016/j.neuron.2020.06.030

Peer, M., Brunec, I. K., Newcombe, N. S., & Epstein, R. A. (2021). Structuring Knowledge with Cognitive Maps and Cognitive Graphs. Trends in Cognitive Sciences, 25(1), 37–54. 10.1016/j.tics.2020.10.004

Pfeiffer, B. E., & Foster, D. J. (2013). Hippocampal place-cell sequences depict future paths to remembered goals. Nature, 497(7447), 74–79. 10.1038/nature12112

Piray, P., & Daw, N. D. (2021). Linear reinforcement learning in planning, grid fields, and cognitive control. Nature Communications, 12(1), 4942. 10.1038/s41467-021-25123-3

Rueckemann, J. W., Sosa, M., Giocomo, L. M., & Buffalo, E. A. (2021). The grid code for ordered experience. Nature Reviews. Neuroscience, 22(10), 637–649. 10.1038/s41583-021-00499-9

Russek, E. M., Momennejad, I., Botvinick, M. M., Gershman, S. J., & Daw, N. D. (2017). Predictive representations can link model-based reinforcement learning to model-free mechanisms. PLoS Computational Biology, 13(9), e1005768. 10.1371/journal.pcbi.1005768

Sanguinetti-Scheck, J. I., & Brecht, M. (2020). Home, head direction stability, and grid cell distortion. Journal of Neurophysiology, 123(4), 1392–1406. 10.1152/jn.00518.2019

Sargolini, F., Fyhn, M., Hafting, T., McNaughton, B. L., Witter, M. P., Moser, M.-B., & Moser, E. I. (2006). Conjunctive representation of position, direction, and velocity in entorhinal cortex. Science (New York, N.Y.), 312(5774), 758–762. 10.1126/science.1125572

Schacter, D. L., Addis, D. R., & Buckner, R. L. (2007). Remembering the past to imagine the future: The prospective brain. Nature Reviews Neuroscience, 8(9), Article 9. 10.1038/nrn2213

Sorscher, B., Mel, G. C., Ocko, S. A., Giocomo, L. M., & Ganguli, S. (2023). A unified theory for the computational and mechanistic origins of grid cells. Neuron, 111(1), 121-137.e13. 10.1016/j.neuron.2022.10.003

Spelke, E. S. (2022). What Babies Know: Core Knowledge and Composition Volume 1. Oxford University Press.

Stachenfeld, K. L., Botvinick, M. M., & Gershman, S. J. (2017). The hippocampus as a predictive map. Nature Neuroscience, 20(11), 1643–1653. 10.1038/nn.4650

Tolman, E. C. (1948). Cognitive maps in rats and men. Psychological Review, 55(4), 189–208.

Tsividis, P. A., Loula, J., Burga, J., Foss, N., Campero, A., Pouncy, T., Gershman, S. J., & Tenenbaum, J. B. (2021). Human-Level Reinforcement Learning through Theory-Based Modeling, Exploration, and Planning (2107.12544). arXiv. 10.48550/arXiv.2107.12544

Tukker, J. J., Beed, P., Brecht, M., Kempter, R., Moser, E. I., & Schmitz, D. (2022). Microcircuits for spatial coding in the medial entorhinal cortex. Physiological Reviews, 102(2), 653–688. 10.1152/physrev.00042.2020

Wernle, T., Waaga, T., Mørreaunet, M., Treves, A., Moser, M.-B., & Moser, E. I. (2018). Integration of grid maps in merged environments. Nature Neuroscience, 21(1), Article 1. 10.1038/s41593-017-0036-6

Whittington, J. C. R., Muller, T. H., Mark, S., Chen, G., Barry, C., Burgess, N., & Behrens, T. E. J. (2020). The Tolman-Eichenbaum Machine: Unifying Space and Relational Memory through Generalization in the Hippocampal Formation. Cell, 183(5), 1249-1263.e23. 10.1016/j.cell.2020.10.024

Widloski, J., & Foster, D. J. (2022). Flexible rerouting of hippocampal replay sequences around changing barriers in the absence of global place field remapping. Neuron, 110(9), 1547-1558.e8. 10.1016/j.neuron.2022.02.002

